# Rhesus macaques as a tractable physiological model of human ageing

**DOI:** 10.1101/2020.06.10.143669

**Authors:** Kenneth L. Chiou, Michael J. Montague, Elisabeth A. Goldman, Marina M. Watowich, Sierra N. Sams, Jeff Song, Julie E. Horvath, Kirstin N. Sterner, Angelina V. Ruiz-Lambides, Melween I. Martínez, James P. Higham, Lauren J. N. Brent, Michael L. Platt, Noah Snyder-Mackler

## Abstract

Research in the basic biology of ageing is increasingly identifying mechanisms and modifiers of ageing in short-lived organisms such as worms and mice. The ultimate goal of such work is to improve human health, particularly in the growing segment of the population surviving into old age. Thus far, few interventions have robustly transcended species boundaries in the laboratory, suggesting that changes in approach are needed to avoid costly failures in translational human research. In this review, we discuss both well-established and alternative model organisms for ageing research and outline how research in nonhuman primates is sorely needed, first, to translate findings from shorter-lived organisms to humans, and second, to understand key aspects of ageing that are unique to primate biology. We focus on rhesus macaques as a particularly promising model organism for ageing research due to their social and physiological similarity to humans as well as the existence of key resources that have been developed for this species. As a case study, we compare gene regulatory signatures of ageing in the peripheral immune system between humans and rhesus macaques from a free-ranging study population in Cayo Santiago. We show that both mRNA expression and DNA methylation signatures of immune ageing are broadly shared between macaques and humans, indicating strong conservation of the trajectory of ageing in the immune system. We conclude with a review of key issues in the biology of ageing for which macaques and other nonhuman primates may uniquely contribute valuable insights, including the effects of social gradients on health and ageing. We anticipate that continuing research in rhesus macaques and other nonhuman primates will play a critical role in conjunction with model organism and human biodemographic research in ultimately improving translational outcomes and extending health and longevity in our ageing population.

## Introduction

The identification and development of biological models of human ageing have been the focus of decades of work. Indeed, research in organisms ranging from yeast to worms to mice has provided exceptional insight into how cellular and molecular processes change with age. From a comparative perspective, this work has provided important insights into the evolution of senescence across the tree of life [1]. From a biomedical perspective, such work has aimed to develop interventions that slow the pace of age-related declines and thus treat or prevent a host of diseases and ailments associated with older individuals—a central goal of geroscience. Short-lived laboratory organisms are limited, however, in their translational impact due to inherent biological dissimilarity stemming from their wide evolutionary distance from humans. In fact, therapeutics developed based on preclinical studies of short-lived laboratory organisms encounter a high failure rate in clinical trials, both in the ageing domain [2] and otherwise [3].

Nonhuman primates (NHPs) are our closest living relatives and are thus natural intermediate models bridging the gap between rodents and clinical human trials. Due to their phylogenetic affinity, NHPs tend to resemble humans in myriad anatomical, physiological, behavioural, and life history traits. Of particular relevance for ageing research, NHPs resemble humans in having extended longevity coupled with exceptionally slow life histories. Primates live approximately twice as long as predicted by their body size [4] and have long juvenile periods and delayed ages of sexual maturity. Nevertheless, all NHP species exhibit short lifespans compared to humans, thus offering distinct advantages over longitudinal research in humans. Longevity in primates also reflects the success of their evolved defences against ageing processes themselves and could help correct a bias in the field towards short-lived species that have demonstrably poor defences against ageing [4].

Here, we review the use of the most commonly studied NHP, rhesus macaques (*Macaca mulatta*), as a model organism for ageing research, focusing on their utility as a translational model for human health. As a case study, we then compare gene expression and DNA methylation data from peripheral blood between humans and macaques and identify parallel gene regulatory signatures of immune ageing. We conclude by discussing how prospective research in rhesus macaques can most effectively maximise progress in the biology of ageing and, in conjunction with other model organism research, yield important and potentially translational insights into ageing in humans.

## Models for human health and ageing

Our knowledge of ageing in eukaryotes is primarily drawn from four model organisms: yeast (*Saccharomyces cerevisiae*), worms (*Caenorhabditis elegans*), flies (*Drosophila melanogaster*), and mice (*Mus musculus*). These model systems offer distinct advantages for laboratory ageing research including short lifespans, ease of care, propensity for genetic or environmental manipulation, availability of inbred and/or engineered genetic strains, compatibility with powerful molecular genetic techniques, and established track records in disease research. Perturbations to certain longevity-related pathways, particularly ones related to nutrient sensing such as insulin/IGF-1 and mTOR signalling, appear to have conserved effects across these distantly related systems, raising hopes for translation to humans [5]. Dietary restriction, accordingly, shows by far the most compelling and robust evidence of anti-ageing effects of any intervention to date, with significant extension of lifespan or healthspan in all species with sufficient data, including NHPs [6,7].

Short lifespans have long been considered an advantage in laboratory ageing research. There is currently substantial interest, for instance, in developing the turquoise killifish (*Nothobranchius furzeri*) as a vertebrate ageing model due to its extremely short lifespan (4–9 months) compared to mice (2–4 years) [8]. Investigation of organisms that exhibit extreme longevity, however, may also hold unique promise for identification of anti-ageing defences obtained through evolution. Our own species is an obvious candidate, and comparison among human populations and among individuals within populations—including, for instance, centenarians—has shed tremendous light on genetic and environmental effects on longevity [9]. Some species of bivalves have extreme lifespans—the ocean quahog (*Arctica islandica*), for instance, is notable for living up to 507 years [10]. The naked mole-rat (*Heterocephalus glaber*) is another potential model, living 5–10 times longer than the similarly sized mouse. They are also devoid of cancer and, unlike all other studied mammals, do not appear to senesce [11].

No other animals share our present environments to the extent that companion animals do, making species such as dogs (*Canis lupus familiaris*) potentially invaluable models of ageing. Dogs also exhibit similar overall patterns of morbidity and comorbidity with age [12]. Given their exceptional genetic and phenotypic diversity, an opportune infrastructure of pet owners and veterinary clinicians, and considerable public interest in extending health and longevity in companion animals, ongoing studies of dog ageing are a promising tool for translational ageing research [13].

Humans are primates, making NHPs obvious candidates that resemble human biology more closely than previously mentioned ageing models. The desire for a small-bodied, relatively short-lived NHP analogue of the mouse model has sparked interest primarily in two species: the grey mouse lemur (*Microcebus murinus*) and the common marmoset (*Callithrix jacchus*). Mouse lemurs and marmosets diverged from humans approximately 74 and 43 million years ago, respectively [14]. Their small sizes (60 g, *M. murinus;* 320 g, *C.jacchus*) short lifespans (5–10 years, *M. murinus;* 5–7 years, *C. jacchus*), short generation times (8–10 months, *M. murinus*; 20 months, *C. jacchus*), and docile temperaments make both species practical and inexpensive to maintain in laboratories, particularly compared to larger-bodied monkeys and apes. The mouse lemur is attracting interest as a biomedical model due to its smaller size, greater fecundity, and faster life history, as well as the presence of parallel age-associated diseases including neurodegenerative pathologies [15]. The marmoset is far more established in laboratory settings, shares a greater phylogenetic relatedness to humans, and exhibits similar age-related diseases including diabetes, cardiovascular disease, and cancer [16]. Both species, however, exhibit peculiarities in their biology that could potentially complicate translation to humans. Mouse lemurs, for instance, seasonally enter a state of daily torpor, while marmosets frequently exhibit reproductive twinning combined with hematopoietic chimerism in their offspring.

Humans are hominoids (apes), making nonhuman apes phylogenetically best-positioned to model human ageing. Chimpanzees (*Pan troglodytes*) and bonobos (*Pan paniscus*) are our closest extant relatives (diverged 6–7 million years ago) [14], and chimpanzees have historically been by far the most commonly encountered ape in biomedical research. Ethical concerns and their endangered status, however, have drastically curtailed their use in laboratory settings in the past decade [17], and many countries have banned invasive great ape research altogether. Field studies, combined with observational and veterinary work in zoos and sanctuaries are generating valuable information on physical, behavioural, and physiological ageing in chimpanzees, with some longitudinal projects even traversing multiple lifespans [e.g., 18]. High costs of care, extremely long lifespans, ethical considerations, and the endangered status of virtually all apes in the wild generally preclude the use of other ape species as viable alternatives to chimpanzee biomedical research.

Given these challenges, cercopithecoids (Old World monkeys), which diverged from humans approximately 29 million years ago [14], serve an invaluable role as the most similar nonhuman proxies of human biology that can be studied in laboratory settings. Apart from macaques, commonly studied examples in laboratory settings include green monkeys (*Chlorocebus sabaeus*) and baboons (*Papio* spp.). Green monkeys, sometimes referred to as vervet monkeys (which more commonly refers to *C. pygerythrus* or all *Chlorocebus*), are a promising model due to their small size and lower cost of care, investments in richly phenotyped captive populations, and the existence of a wild population in the Caribbean descended from an anthropomorphic bottleneck [19]. Baboons are valuable models for genetic studies of complex diseases in large part because of the depth of genotyping and phenotyping efforts in pedigreed populations [20] and the breadth and longevity of longitudinal studies in wild populations [21].

## A rhesus macaque model of health and ageing

Rhesus macaques (*Macaca mulatta*) are the most common NHP in captivity and have been a useful model for human ageing for decades [22–25]. As a major biomedical model, rhesus macaques are the beneficiaries of investment in major resources including high-quality genome assemblies [24,26] and neuroanatomical atlases [27]. The rhesus macaque lifespan is approximately 3–4 times shorter than that of humans (figure 1), yet within this compressed period ageing macaques exhibit a host of parallel declines in physical health, physiological integrity, and brain function [23].

**Figure 1.**
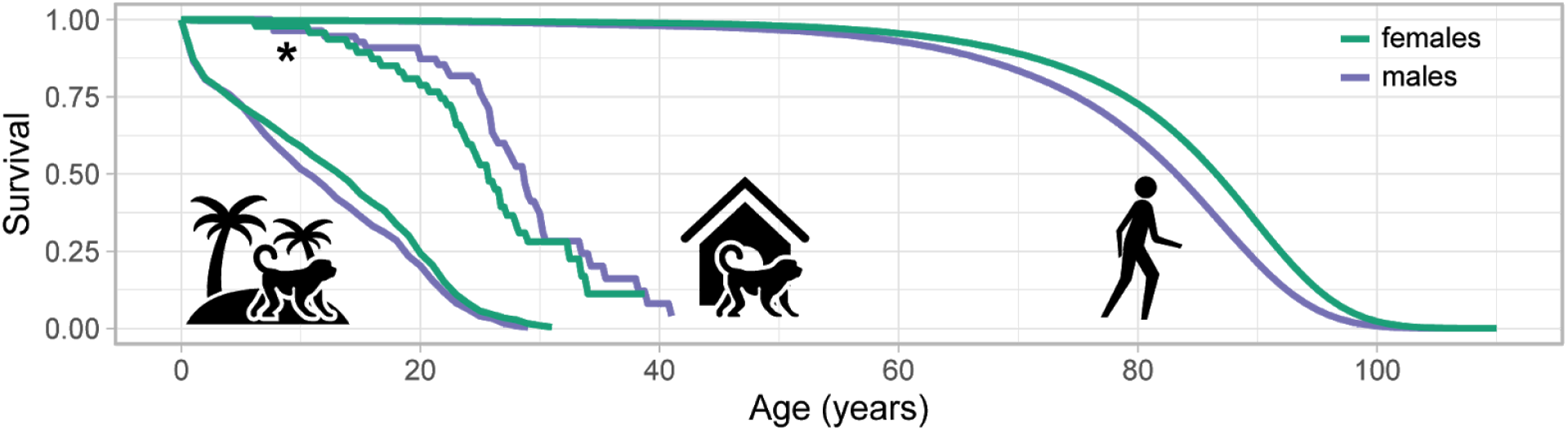
Comparison of survival between rhesus macaques (free-ranging and captive) and humans. Survival curves for free-ranging macaques are from the island population of Cayo Santiago and include 11,659 monkeys born since 1938. Survival curves for captive macaques are from a previously published caloric restriction study [6] (data kindly shared by Rozalyn Anderson, Ricki Colman, and Julie Mattison) and include 102 monkeys fed a control diet across two indoor-housed colonies at the University of Wisconsin–Madison and the National Institute on Aging. These curves (indicated by an asterisk) should be interpreted with caution as monkeys were enrolled at ages ranging from 1–23, introducing selection bias and removing sources of infant mortality from the curve. The true median lifespan is therefore lower than indicated, though still higher than that of free-ranging macaques. Kaplan-Meier curves were estimated for both macaque datasets on right-censored data. Survival curves for humans were plotted from life tables (10-year periods) for the Swedish population obtained from the Human Mortality Database (https://mortality.org) and filtered to the period 2010–2018.

In the wild, rhesus macaques inhabit the largest geographic range of any NHP species and are highly adaptable to human environments, where they are commonly encountered in urban areas. Rhesus macaques are one of 23 species from the macaque genus [28] and in captivity are subdivided into two genetically distinct populations or subspecies, usually referred to as Indian-origin and Chinese-origin rhesus macaques. Several other macaques are also important biomedical species, particularly cynomolgus (also known as long-tailed or crab-eating) macaques (*M. fascicularis*) [29]. The abundance of multiple laboratory macaque species or subspecies, sometimes housed in the same research facilities, creates opportunities for identifying conserved pathways and validating translational interventions across a smaller evolutionary time scale relative to humans and existing model organisms. Validation across distinct macaque taxa could thus function as an important benchmark prior to clinical testing in humans.

Along with all Old World monkeys, rhesus macaques are our closest extant non-ape relatives and thus share many aspects of our anatomy, physiology, neurobiology, and behaviour. Rhesus macaques live approximately 25 years in captivity, with a maximum recorded lifespan of 40 years. While compressed by comparison, the relative timing of their life history parallels that of humans, from development through maturation, reproduction, and senescence. Moreover, the signs and symptoms of ageing manifest in similar ways. For instance, in captivity, rhesus macaques show signs of thinning and greying hair, atrophying skin, and declining motor activity [23,25]. They also naturally exhibit pathological signs of a variety of age-associated diseases—including cardiovascular disease, cancer, and altered glucose metabolism [6,22]—making rhesus macaques valuable disease models for chronic conditions that cannot be adequately studied in short-lived organisms.

Rhesus macaques undergo structural changes to their musculoskeletal biology that contribute to their increasing frailty with age. Apart from decreases in stature [30], ageing rhesus macaques show decreases in bone mineral density and content [31], muscle mass and function [32], and deterioration of spinal cartilage [33]. Rhesus macaques also exhibit age-related changes in body composition, including a redistribution of body fat and increase in overall adiposity [6].

In the brain, ageing rhesus macaques undergo a number of structural changes, such as an increase in microglia density [34] and a loss of synapses and dendritic spines in some cortical areas [35], which may be related to observed impairments in behavioural, sensory, and cognitive functions [35,36]. Transcriptomic and proteomic studies are increasingly identifying age-associated gene and protein expression changes in a variety of brain regions [37–39]. Notably, while rhesus macaque brains exhibit an increased frequency of amyloid plaques with age, a classic hallmark of Alzheimer’s disease, the plaques are not correlated with cognitive decline [35]. Moreover, the increases in amyloid are not characterised by increases of the *Aβ42* form of the protein as in individuals with Alzheimer’s disease. This, combined with the scarcity of neurofibrillary tangles [but see 40], suggests that rhesus macaques do not recapitulate all of the pathologies of Alzheimer’s disease.

### Natural laboratories for ageing research

Laboratory organism research generates tremendous insight into ageing biology. Inherent to the success of laboratory research, however, is the need to reduce natural complexity in exchange for increased experimental control. While undoubtedly a necessary tradeoff, it creates a problem in that modern humans, the ultimate target of biomedical research, are a genetically structured and diverse population living in highly variable and complex ecological and sociocultural environments.

NHPs, and rhesus macaques specifically, are valuable as models due to the impressive breadth and depth of efforts to understand species in the wild [41] or in naturalistic environments [42]. While experimental design is necessarily constrained in such settings, field studies play at least one crucial role for ageing research in that they reveal heterogeneity in ageing under environmentally realistic circumstances. By leveraging natural genetic and environmental variation, and by using research tools such as noninvasive biomarkers or capture-and-release sampling, field research is well-positioned to monitor how genes and the environment interact to influence the ageing trajectory. While these approaches are sometimes handicapped in their statistical power relative to laboratory organism research, they are similar to methods used by human biodemographers and function as an important complement to laboratory research.

The population of rhesus macaques on the island of Cayo Santiago has served as a unique resource for macaque research for over 80 years [43]. Maintained by the Caribbean Primate Center at the University of Puerto Rico, the population was founded in the late 1930s, during which 409 founders were imported from India and released onto the 15.2-hectare island located 1 km from the eastern coast of Puerto Rico. Since then, the macaques have flourished, with the population now approaching 2,000 individuals. While the macaques are provisioned with commercial feed, tattooed for identification, and subject to periodic capture-release, they otherwise live under naturalistic circumstances without human interference. Access to feed, for instance, is subject to social competition [44].

These properties provide a unique balance of the costs and benefits of laboratory and wild macaque research. Consequently, the population has been subject to intensive study, including in the context of ageing and senescence [45–47]. Because the population has been studied continuously since its founding (with constant surveillance commencing in 1956), all individuals on the island are identified and matrilines are known stretching back to the colony’s founding. Full pedigrees are known stretching back to the 1980s with the advent of genetic paternity testing. Importantly, animals are free to form their own social associations, and organise into the multimale-multifemale social structure seen in wild rhesus macaques. The Cayo Santiago macaques are free of natural predators, but without housing and with minimal veterinary intervention, they are subject to all environmental stresses including parasites and pathogens. Consequently, the Cayo Santiago macaques have a shorter lifespan than captive macaques, with females living to a median of 18 years and a maximum of 31 years [42] (figure 1). The combination of geographic confinement on an isolated island setting with periodic capture-and-release efforts that enable monitoring of critical biomarkers of ageing ensures that the Cayo Santiago population will continue to serve as a vital resource for ageing and other biomedical research.

### Parallel immune signatures of ageing in rhesus macaques and humans

To demonstrate the value of rhesus macaques as an ageing model, we compared signatures of ageing in rhesus macaque and human ageing in the peripheral immune system using data obtained from the Cayo Santiago population. We focus on the peripheral immune system because of its convenience—whole blood is a readily available biological sample, particularly in larger-bodied animals—and, critically, because the immune system exhibits numerous well-documented changes with age.

The immune system is vital for responding to the threat of foreign pathogens. It is widely accepted that the immune system undergoes numerous changes in aged individuals, leading to a state of immunosenescence [48,49]. One of the strongest hallmarks of ageing in peripheral blood is a reduction in the number of naïve T cells and an accumulation of late-stage differentiated memory T cells [50]. Another hallmark of ageing is a steady increase in both pro- and anti-inflammatory activity in innate immune cells, particularly macrophages and monocytes. This increase in inflammatory activity is thought to be linked to a state of chronic, low-grade inflammation that characterises ageing, a phenomenon that has been termed inflamm-ageing [51].

Like humans, rhesus macaques demonstrate marked changes in immune function with age, including the expected reduction in undifferentiated T cells and increase in differentiated memory T cells [52,53]. Evidence for increases in circulating proinflammatory cytokines has been more mixed [52], although some recent evidence suggests increases in some cytokines, including IL-6 [53, 54, but see 55]. In innate immune cells, however, ageing has mixed effects on responses to the binding of pro-inflammatory cytokines, with most responses declining with age [54].

Broadly, age-related changes in macaque peripheral immune function largely parallel those in humans. Direct comparisons between macaques and humans, however, have thus far been largely qualitative, with a focus on limited numbers of key biomarkers. Here, we aimed to empirically estimate similarity in age-related changes in peripheral immune function between humans and rhesus macaques. We quantified age associated changes in immune cell function by assaying blood gene expression and CpG methylation in macaques and compared our data to published data from human populations.

We sequenced whole-blood transcriptomes using RNA-seq from a total of 48 macaques spanning the natural age distribution of Cayo Santiago (figure S1a). After mapping and filtering (SI methods), we modelled the effects of chronological age—controlling for sex as a covariate—on mRNA expression across 11,150 genes using mixed models to correct for the kinship structure in our dataset [56,57]. We found that age-associated increases in gene expression were enriched for biological processes including inflammatory responses and responses to virus (table S1), while age-associated decreases in gene expression were enriched for biological processes including recognition processes in immunoglobulin production, phagocytosis, and B cell receptor signalling (table S2). These results are consistent with previous findings that ageing is associated with increased inflammatory responses, declining B cell numbers and functions, and impaired phagocytosis in macrophages [48].

We then compared our results to a published set of 1,497 age-associated genes from an analysis of ~15,000 humans of European ancestry [58]. We were interested in whether genes exhibit similar age-related changes in expression between macaques and humans; hence we evaluated for each gene whether macaques and humans showed equivalent directions of change in gene expression. We were able to evaluate a total of 970 genes measured in both datasets. After filtering to include only the 70 genes passing a false discovery rate (FDR) threshold of 20% in both datasets—with the relatively low fraction likely explained by greater power in the human dataset—we found that 67 (95.7%) were concordant, indicating that their expression changed with age in the same direction (figure 2a). 3 genes that were discordant were involved in functions such as lipid metabolism (*DGKG, LPCAT1*) and endoplasmic reticulum/Golgi apparatus transport (*ERGIC1*). This similarity decreased as we relaxed our significance threshold for inclusion of genes, but even when implementing the most permissive threshold (all 970 genes), we still found that 74.2% of genes were consistent in direction. This indicates that the magnitude of similarity in the directionality of peripheral blood immune ageing is sufficiently strong so as to compensate for increased statistical noise in our results, and that increasing our sample size and statistical power would increase both the number and fraction of genes consistent in direction.

**Figure 2.**
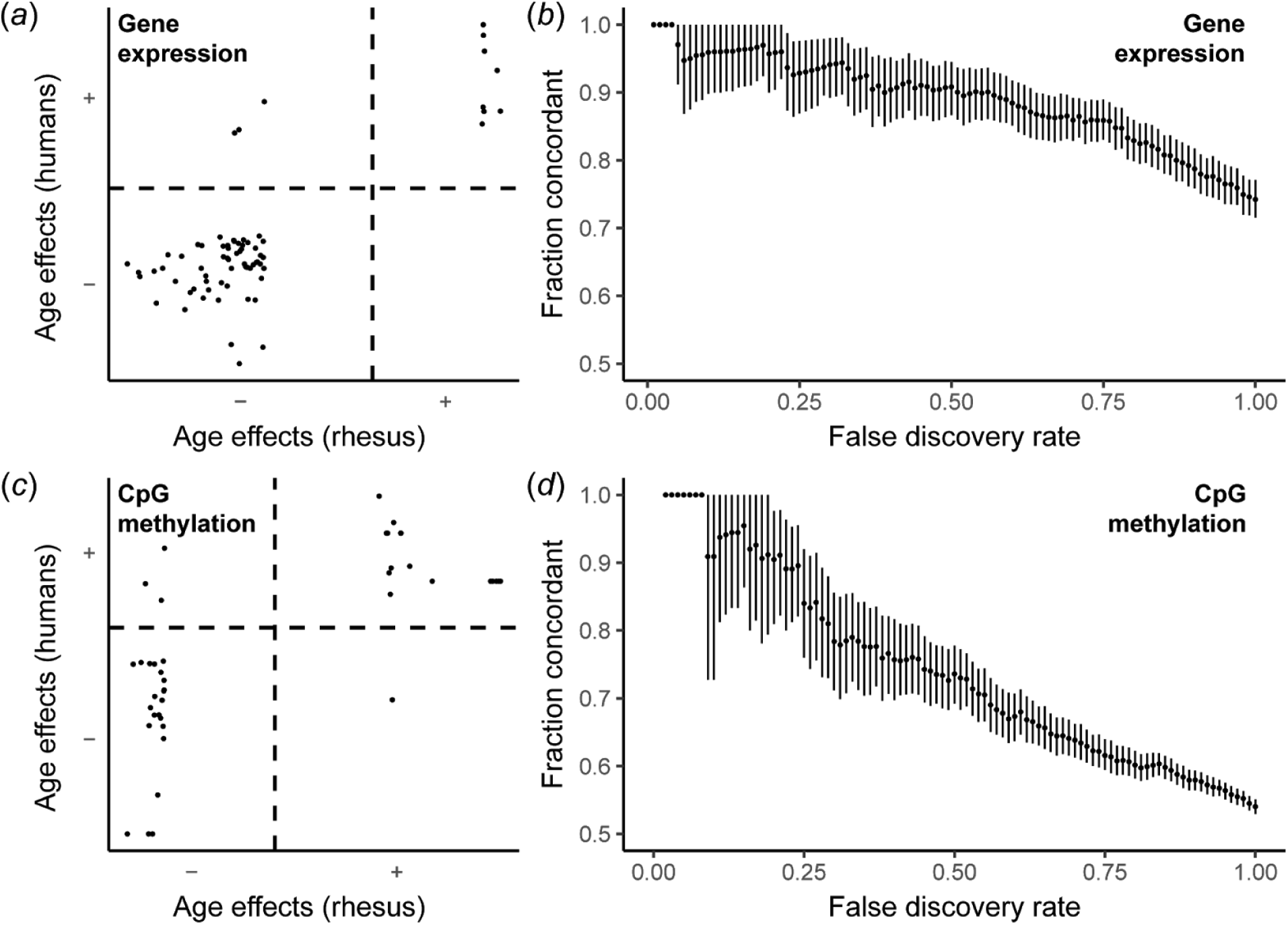
Comparisons of ageing effects in peripheral blood between macaques and humans. In these plots, (a) Comparison of age effects in gene expression. Standardised *β* values from our rhesus RNA-seq analysis are plotted against *Z*-scores in single-copy orthologous genes from a published analysis of humans [58]. Genes not passing a false discovery rate threshold ≤20% for both datasets are not shown. Out of 70 genes passing this threshold, 67 (95.7%) were consistent in sign, indicating shared directional changes with age. (b) Comparison of concordance in direction of age effects in gene expression and its sensitivity to statistical power. The false discovery rate threshold was adjusted across a range of values and concordance was estimated as the proportion of genes with the same sign after filtering. (c) Comparison of age effects in CpG methylation. Standardised *β* values from our rhesus RRBS analysis are plotted against standardised *β* values of orthologous CpGs from our analysis of a published methylation array dataset of 656 humans [59]. CpGs not passing a false discovery rate threshold ≤20% for either dataset are not shown. Out of 42 CpGs passing this threshold, 38 (90.4%) were consistent in sign, indicating shared directional changes with age. (d) Comparison of concordance in direction of age effects in CpG methylation and its sensitivity to statistical power. The false discovery rate threshold was adjusted across a range of values and concordance was estimated as the proportion of genes with the same sign after filtering.

Next, we tested if phylogenetic conservatism was such that age prediction models developed using human transcriptomes could accurately predict chronological ages in macaques. To test this, we used a transcriptomic age prediction approach developed in humans [58] to predict chronological ages from our normalised macaque gene expression data. We included genes that passed an FDR of 20% in both analyses. Following previous studies [58], predictions were agnostic to sex as sex differences in perpheral blood transcriptomic ageing tend to be relatively minor. After scaling the predictions to the age distribution in our sample population, we found a strong correlation between known chronological ages and cross-species transcriptomic age predictions, with a Pearson’s *r* of 0.681 (*p* = 1.0E-7). These results in fact fell within the range of variation in model predictions across human cohorts, with Pearson’s *r* ranging from 0.320 to 0.774 [58]. The average difference between predicted and known ages was also comparable, with our mean difference (3.22 years) falling in the upper range of values across human cohorts (4.84 – 11.21 years) [58] after taking into account differences in lifespan. This mean difference increased with age—for instance, average error in age predictions was 5.38 years for animals over 15 years old—suggesting that human predictors may not capture ageing trajectories in older macaques or that there may be greater heterogeneity in cumulative ageing outcomes among older surviving free-ranging macaques.

**Figure 3.**
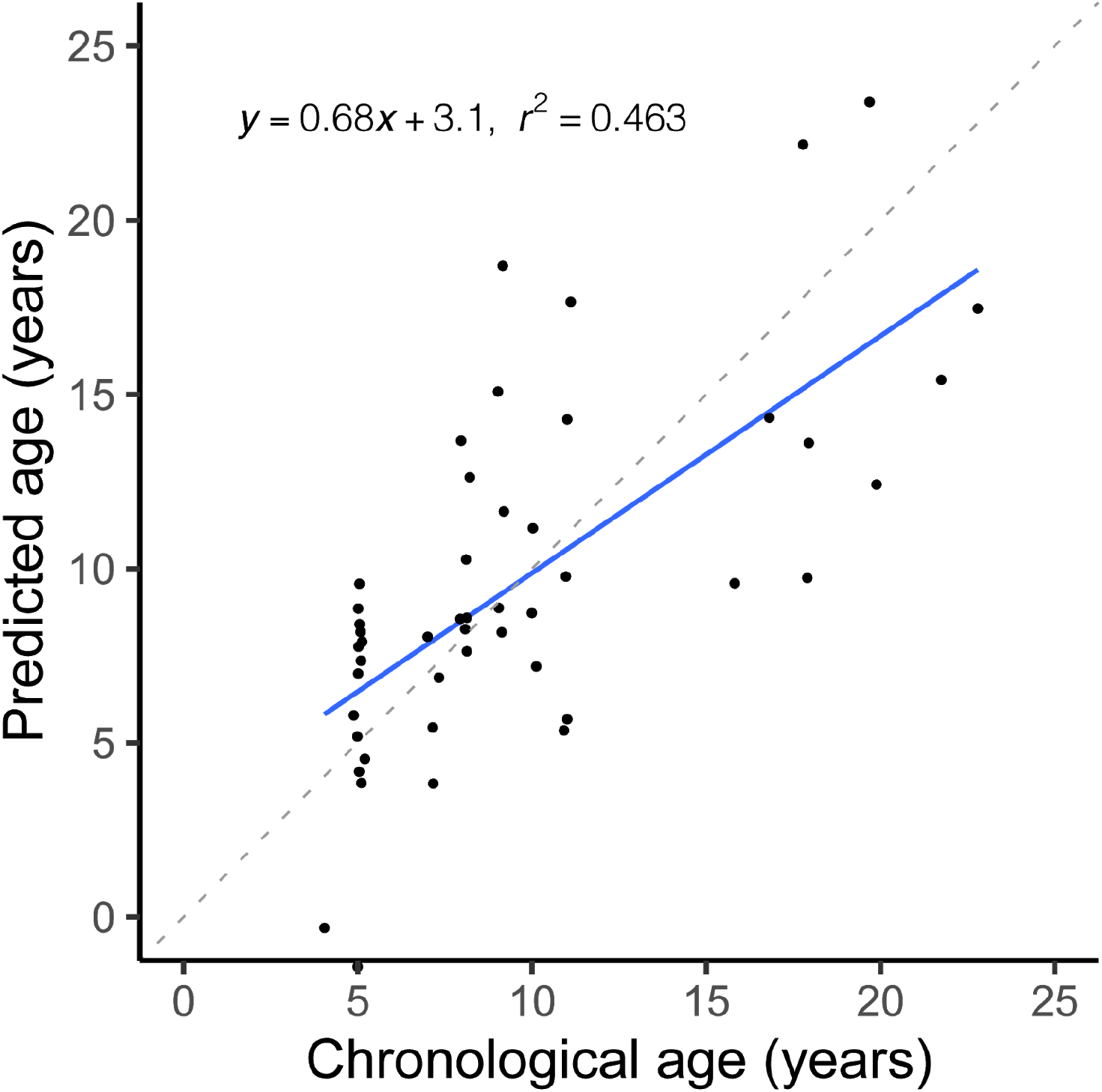
Interspecies transcriptomic predictions of rhesus macaque ages using a transcriptomic clock developed in humans [58] show a strong correlation with known chronological ages. The regression line is shown in blue while the expected relationship (*y* = *x*) is shown as a dashed line in light grey.

We also evaluated the similarity in gene regulatory changes with age between macaques and humans. We used CpG methylation because it is a key epigenetic regulator of transcription, one of the best-characterised forms of gene regulation, and exhibits highly predictable changes with age in humans [59,60]. We assayed CpG methylation in 104 macaques spanning the natural age distribution (figure S1b) using reduced-representation bisulfite sequencing (RRBS) [61]. After filtering (see SI methods), we modelled methylation changes across 40,726 CpGs as a function of age—including sex as a covariate—using a binomial mixed modelling approach that controls for population structure [62].

Similar to our gene expression analysis, we compared our age effects for all CpG sites against orthologous sites or regions in a published methylation array dataset in humans [59]. Because of limited overlap between CpG sites represented in the array and RRBS datasets, and because DNA methylation patterns are highly correlated at neighbouring sites within 1–2 kb of one another [63], we considered CpG sites in the human array orthologous if they were the closest available site within 100 bp of a macaque CpG site. We thus compared 8,482 sites (regions) in this manner.

Among 42 sites that passed a significance cutoff (FDR ≤ 0.2) in both datasets, we found that 38 sites (90.5%) were concordant in sign (figure 2c). When we evaluated the concordance in the directionality of age effects across a range of FDR thresholds, we similarly found lower proportions of concordant sites across all significance thresholds. Even at the most permissive threshold (FDR ≤ 1.0), we found that 54% of sites were concordant in direction. These results were largely robust to our choice of criteria for identifying orthologous sites (figure S2). Due to limited overlap with well-established epigenomic-clock CpGs in humans [59,60], we did not attempt to predict macaque chronological ages using our methylation dataset.

Our results as a whole reveal extensive similarities in transcriptomic and epigenomic signatures of peripheral immune ageing between rhesus macaques and humans. This effect was strongest at an FDR threshold < 20%, where the confidence intervals encompassed 100% concordance for both the CpG methylation and gene expression comparisons (figure 2b, 2d). For both analyses, after filtering out non-single-copy orthologs and non-significant effects, we found that a relatively small number of genes or sites remained. This is in large part a reflection of lower statistical power in our rhesus macaque dataset, which causes a loss of non-significant genes/sites in our dataset that could otherwise be compared to genes/sites in the more highly powered human datasets.

Despite some caveats, the analyses here broadly demonstrate that age-related changes in immune function are qualitatively similar between humans and rhesus macaques and lay the foundation for future work quantifying similarities in immune ageing.

## Outlook and challenges

### Ageing across the body

The human peripheral immune system is a frequent target of study for many reasons, not least of which is that whole blood can be sampled relatively noninvasively, repeatedly, and in large quantities from volunteers. Ageing in the peripheral immune system, however, is part of a multifaceted process involving myriad interdependent physiological systems and tissues. A challenge for future geroscience research is to integrate data across a wide range of physiological systems into a broad and generalisable model of human ageing [60].

Ethical and logistical considerations restrict the ability to assess ageing in humans in most tissue types apart from blood. Here, rhesus macaques provide tremendous value by way of their shared biology, particularly for organs that often require post-mortem sampling such as the heart and brain. Cross-sectional studies of such rare samples promise to shed valuable light on poorly characterised processes of ageing in such tissue with profound implications for health and disease.

### Heterogeneity and modifiers of the ageing process

Not all individuals age at the same pace and in the same way. Quantifying heterogeneity in ageing and identifying the mechanisms underlying individual resilience to ageing across populations may lead to novel interventions in humans. Pursuing this goal requires insights from both laboratory settings, where important variables can be controlled or manipulated, and field settings, where existing genetic and environmental variation can be leveraged. Rhesus macaque research is well-established in both settings and represents a promising avenue to obtaining a more comprehensive understanding of putative modifiers of ageing, including their mechanisms of action, effects on lifespan, and robustness to variation in genetics and environment.

Diet is a major factor influencing health and lifespan and has correspondingly received substantial attention in the ageing literature. Dietary restriction—particularly caloric restriction—has a robust impact in a variety of organisms, attenuating age-associated declines and extending lifespan in most studied species to date. After initial mixed results, it is now generally accepted that caloric restriction extends both healthspan and lifespan in rhesus macaques [6], establishing rhesus macaques as an important NHP model for the effects of diet on ageing. A key remaining challenge for geroscience is understanding the mechanisms underlying dietary restriction on ageing.

Sex differences in ageing also represent a major gap in knowledge that can be exploited to disentangle intrinsic and extrinsic mechanisms underlying survival and longevity [64]. In humans, a robust female survival advantage is well-documented, although paradoxically, their health late in life appears to be poorer compared to men. The reasons underlying this paradox are currently unknown, although proposed explanations focus on sex differences in extrinsic mortality risk and sex differences in endocrine function [64]. Like humans, free-ranging rhesus macaques appear to show a slight female survival advantage [65] (figure 1). Furthermore, captive macaques under caloric restriction exhibit sex-dependent changes in body composition and nutrient-dependent glucose metabolism [6]. These data also suggest that rhesus macaques may be an opportune model for disentangling sex-dependent effects of extrinsic and intrinsic factors on lifespan. Indeed, male and female macaque mortality varies across the year, which reflects sex differences in the pressures that may affect mortality. Male mortality is highest in the mating season, while female mortality peaks during the birth season [45].

The relationship between the social environment and ageing is emerging as an important area of study. In humans, low socioeconomic status and increased social isolation are major epidemiological risk factors for mortality, mental conditions, and chronic diseases [66,67]. These and other components of social adversity are thought to accelerate ageing through altered hypothalamic–pituitary–adrenal (HPA) axis function, particularly a gradual desensitisation of the glucocorticoid receptor response, leading to chronic activation of downstream proinflammatory cascades [68]. Indeed, various forms of social adversity are associated with elevated expression of proinflammatory genes and decreased expression of genes related to innate immune responses in humans [69] and rhesus macaques [70]. Low macaque dominance rank, for instance, has been causally linked to altered glucocorticoid and immune regulation and a polarisation of some immune pathways towards a proinflammatory response [71–74]. Immune signatures of social status in macaques and ageing in humans have also been shown to overlap substantially in their transcriptional profiles [75], suggesting that chronic social adversity may accelerate physiological aging. Further supporting this idea, social integration—a condition that is believed to buffer against adverse effects of social stress—predicts survival in female macaques [42,76].

Studies of social environmental effects on ageing must also disentangle confounded relationships between social and asocial factors. Low socioeconomic status, for instance, is associated with poor nutrition and reduced access to healthcare [67]. These issues can be mitigated in animal models, for which complexities of the social and physical environment can be standardised. Compared to other organisms such as rodents, macaques are a particularly promising model because their expressed social structure—large multimale-multifemale groups with gradients in social hierarchy and social integration—resembles our own. In order to identify human-relevant modifiers of ageing, it is therefore crucial to study the aetiology of ageing and associated conditions in socially and ecologically relevant organisms and settings.

## Conclusion

Rhesus macaques fulfil an important role for ageing research by providing an auspicious balance between the benefits and drawbacks of research in short-lived organisms or in humans. Future innovations in ageing research will undoubtedly be based on cutting-edge techniques developed for short-lived laboratory model organisms as well as insights gleaned from human research into aspects of biology or disease that do not reproduce well in macaques.

Humans, the ultimate target of translational ageing research, are an evolutionarily unique, genetically heterogeneous (with some structure) population living in extremely variable environments and with tremendous sociocultural variation. Existing model species almost certainly do not fully capture our biology and diversity, which likely explains the high failure rate of translational clinical trials. Our view is therefore that geroscience is best served with a greater diversity of model organisms. Preclinical research that concludes with mice requires a leap across nearly 100 million years of independent evolution. A more manageable strategy may be to first replicate mouse-based findings in more closely related taxa such as rats. Likewise in NHPs, strategic investments in multiple viable models including mouse lemurs, marmosets, baboons, and multiple species of macaque may afford opportunities for understanding conserved hallmarks of ageing across primates, increasing the probability of success in human trials, which are costly both in time and expense.

## Data accessibility

Raw reads are deposited in the NCBI Sequence Read Archive under BioProject accession number PRJNA610241. Sample metadata and analysis code are available on GitHub and archived on Zenodo (doi:10.5281/zenodo.3747243).

## Author contributions

K.L.C., J.E.H., J.P.H., L.J.N.B., M.L.P. and N.S.M. conceived the research. K.L.C., M.J.M., J.E.H., K.N.S., A.V.R.L., M.I.M., J.P.H., L.J.N.B., M.L.P., and N.S.M. designed the study. K.L.C., M.J.M., E.A.G., M.W., S.N.S., J.S., and N.S.M. performed the research. K.L.C. and N.S.M. wrote the paper. All authors read and approved the final manuscript.

## Acknowledgements

We are grateful to the Cayo Santiago census team—Giselle Caraballo, Nahiri Rivera, Bianca Giura, and Julio Resto—and caretakers; to Rozalyn Anderson, Ricki Colman, Julie Mattison, and Raisa Hernández Pacheco for sharing survival data; and to Melissa Emery Thompson and Alexandra Rosati for organizing the 2019 American Association of Physical Anthropologists symposium that gave rise to this paper and special issue.

This research was supported by the National Institutes of Health (NIA R00-AG051764, NCRR/ORIP P40-OD012217, NIA R01-AG060931, NIMH R01-MH096875, NIMH R01-MH089484, NIMH R01-MH118203). K.L.C. is supported by an NIH fellowship (NIA T32-AG000057).

## Ethics statement

All work with animals was reviewed and approved by the Institutional Animal Care and Use Committees of the University of Washington (assurance number A3464-01) and the University of Puerto Rico, Medical Sciences Campus (assurance number A400117). This research adheres to the American Society of Primatologists Principles for the Ethical Treatment of Nonhuman Primates.

