## Supplementary Information for "Rhesus macaques as a tractable physiological model of human ageing"

Electronic Supplementary Material

### Appendix S1. Supplementary methods

##### Sample collection and nucleic acid extraction

Whole blood was drawn from sedated rhesus macaques by veterinary staff as part of routine capture-and-release efforts on Cayo Santiago. Blood was drawn into PAXgene blood RNA tubes (PreAnalytiX GmbH) for RNA analysis and K3 EDTA vacutainers (BD Biosciences) for DNA analysis. All samples were stored at -80°C within 8 hours of collection.

We extracted total RNA from PAXgene-stabilised tubes using the MagMAX for Stabilized Blood Tubes RNA Isolation Kit (ThermoFisher), following manufacturer instructions for maximising RNA quality and yield. We extracted total DNA from whole blood using the DNeasy Blood & Tissue kit (QIAgen) according to manufacturer instructions for Purification of Total DNA from Animal Blood or Cells.

Because PAXgene and EDTA tubes were not always available from the same animals or blood draws, we did not attempt in this study to sequence the same individuals and time points for both RNA and DNA analysis. We instead included individuals spanning the age distribution based on the availability of suitable samples separately for each analysis.

##### Library preparation and sequencing

We measured gene expression using whole-transcript RNA-seq. We enriched mRNA using the NEBNext Poly(A) mRNA Magnetic Isolation Module (New England Biolabs) and prepared cDNA libraries using the NEBNext Ultra II RNA Library Prep Kit for Illumina (New England Biolabs). To prepare the libraries, we used an input RNA quantity of 200 ng, targeted a insert size of 600 bp, and amplified the library with12 PCR cycles. All other procedures followed manufacturer recommendations.

We measured CpG methylation using reduced representation bisulfite sequencing (RRBS). RRBS is effective when existing methylation arrays for a taxon of interest do not exist and is more cost-effective than whole-genome bisulfite sequencing. We prepared RRBS libraries based on previous protocols [1,2] (detailed protocol can be found here: https://smack-lab.com/protocols/). Briefly, 300 ng of DNA was first digested, along with a small amount (0.1 ng) of lambda phage DNA, using *Msp*I (New England Biolabs) to produce fragments with CpG ends. Following digestion, we performed end repair and adapter ligation on *Msp*I-digested DNA using NEBNext Ultra II (New England Biolabs) reagents. We then performed bisulfite conversion using the EZ-96 DNA Methylation-Lightning MagPrep kit (Zymo Research). Libraries were then PCR-amplified for 16 cycles with unique dual indexed sequencing primers. Unless otherwise stated, all procedures followed manufacturer recommendations.

For both RNA-seq and RRBS libraries, libraries were pooled in 1 μl volumes and sequenced on an Illumina MiSeq using the MiSeq v2 Nano kit and 2×150 bp sequencing. Based on proportional read representations, libraries were then re-pooled in equimolar quantities. RNA-seq libraries were sequenced on two Illumina NextSeq 500 flow cells using 2×38 bp sequencing. RRBS libraries were sequenced on an Illumina NovaSeq S2 flow cell using 2×51 bp sequencing.

##### *Gene expression preprocessing and modelling*

We mapped cDNA reads to the rhesus macaque reference assembly (Mmul_8.0.1) using kallisto [3]. We removed lowly expressed genes by filtering out genes with fewer than 2 transcripts per million (TPM) and normalised transcript abundances using voom from the limma package in R for subsequent analysis.

We modelled age effects on expression using an efficient mixed model association (EMMA) test [4] implemented in the EMMREML package in R [5]. EMMA requires a kinship matrix, which it uses to correct for the population structure in the dataset. We used the kin function from the synbreed package [6] in R to generate a kinship matrix using existing pedigree information from Cayo Santiago as input. We then used the emmreml function to model expression against age, including sex and library batch as covariates.

##### *CpG methylation preprocessing and modelling*

After trimming RRBS reads using Trim Galore!, we mapped reads to the rhesus macaque reference assembly (Mmul_8.0.1) and extracted methylated and total read counts using Bismark [7]. We filtered out constitutively hypermethylated and hypomethylated CpGs from our dataset by removing sites with median methylated fraction less than 10% or greater than 90%.

We modelled age effects on CpG methylation using a mixed model association for count data via data augmentation (MACAU) test in the PQLseq package in R [8]. We used the kin function from the synbreed package [6] in R to generate a kinship matrix using existing pedigree information from Cayo Santiago as input. We then used the pqlseq function to model methylated counts as a function of age, while controlling for sex and library preparation data (a potential batch effect).

##### *Enrichment analysis*

For our gene expression analysis, we conducted Gene Ontology (GO) enrichment analyses using topGO [9] to identify pathways that were nonrandomly associated with directional changes in gene expression with age. We restricted our analyses to GO biological processes to focus on pathways with clear links to immune function. We used the "weight01" algorithm to conduct a Kolmogorov–Smirnov test to identify pathways with strongest associations with increased expression with age (positive standardised $\beta$) or decreased expression with age (negative standardised $\beta$). We report GO terms passing a false discovery rate (FDR) threshold of 10%.

##### *Comparison to published human gene expression datasets*

In order to compare our gene expression analysis to analyses of human ageing, we obtained summary files of gene-by-gene effect sizes (*Z* values) and *p* values from Peters *et al*. [10] (“Supplementary Data 1” of the external paper). We compared our results to the 1,497 genes identified in their metaanalysis of a discovery and replication dataset as showing the strongest expression changes with age. We matched genes in the human dataset to rhesus macaque genes using Ensembl homolog annotations obtained via biomaRt [11].

We then compared the concordance of effect sizes for the 970 genes for which we had estimated age effects in both our rhesus macaque dataset and the published human data. We calculated concordance across a range of significance thresholds (FDR from 0 to 1 in 0.01 increments), with bootstrapping (random resampling with replacement) to obtain 95% confidence intervals.

We initially compared our data to the metaanalysis reported by Peters *et al*. [10] because these results incorporated an independent replication dataset and thus produced more strongly supported markers. This set of markers, however, was limited in numbers. To conduct a more comparable comparison, we also compared our data to the discovery results. For this analysis, a total of 7,074 were found in both datasets. With this set, we repeated our analysis of concordance as described above (figure S3).

##### *Prediction of transcriptomic age*

We use the transcriptomic age predictor and equation provided by Peters *et al*. [10] ("Supplementary Data 5" of the external paper) to predict chronological ages in our rhesus macaque dataset from their gene expression data. We estimated the predictor (*Z*) using equation 12 (reproduced below) on our normalised gene expression matrix.

$$Z = \sum_i x_{v(i)} \widehat{b}_{R(i)}$$

We then scaled the predictor to the mean and standard deviation of our rhesus macaque dataset using equation 13 (reproduced below) to obtain estimated ages.

$$SZ = \mu_{age} + (Z - \mu_Z) \times \frac{\sigma_{age}}{\sigma_Z}$$

##### *Comparison to published human CpG methylation datasets*

In order to compare our CpG methylation analyses to analyses of human ageing, we obtained methylation array data generated using the Illumina HumanMethylation450 BeadChip and reported by Hannum *et al.* [12]. We filtered out constitutively hypermethylated and hypomethylated CpGs from the dataset by removing sites with median methylated intensity ratios ($\beta$) less than 10% or greater than 90%. We then used a linear model to regress age against methylated intensity ratios, with plate, sex, and race included as covariates.

In order to match CpG sites between our rhesus macaque results and the human results, we used UCSC liftOver to translate coordinates between the rhesus macaque data mapped to the Mmul_8.0.1 reference assembly and the human array data based on the GRCh36 (hg18) reference assembly. We conducted a series of reciprocal conversions—we used the GRCh37 (hg19) assembly as an intermediary due to the lack of a chain file directly linking the two assemblies—and removed missing or duplicate sites in order to ensure that all remaining sites were single-copy orthologs.

Because of the low number of CpG sites (278) overlapping between datasets after these procedures, we expanded our criteria for matching sites by including neighboring sites. We leveraged the fact that DNA methylation patterns are highly correlated at neighboring sites within 1–2 kb of one another [13], and implemented a relatively conservative criterion by matching CpGs sites in the rhesus macaque dataset to human CpGs only if they were the nearest neighboring site within 100 bp. After matching sites in this manner, we calculated concordance in direction by calculating the absolute value of the effect size in macaques (standardised $\beta$) multiplied by the effect size in humans (standardised $\beta$) and classifying the results as positive or negative. We then adjusted our criterion for determining orthologous sites by comparing results linking neighboring sites within 100 bp, 50 bp, and 0 bp (overlapping) (figure S2). For these analyses, we calculated concordance across a range of significance thresholds (FDR from 0 to 1 in

0.01 increments), with bootstrapping (random resampling with replacement) to obtain 95% confidence intervals.

### Supplementary tables

**Table S1**. Gene Ontology terms enriched for increased gene expression with age (FDR ≤ 0.1).

| GO ID | GO term | FDR-adjusted *p* value |
|---|---|---:|
| GO:0006954 | inflammatory response | 0.015 |
| GO:0007189 | adenylate cyclase-activating G protein-coupled receptor signaling pathway | 0.015 |
| GO:0051607 | defense response to virus | 0.020 |
| GO:0071346 | cellular response to interferon-gamma | 0.037 |
| GO:0006508 | proteolysis | 0.076 |
| GO:0045429 | positive regulation of nitric oxide biosynthetic process | 0.094 |

**Table S2**. Gene Ontology terms enriched for decreased gene expression with age (FDR ≤ 0.1).

| GO ID | GO term | FDR-adjusted *p* value |
|---|---|---:|
| GO:0006412 | translation | < 0.001 |
| GO:0002377 | immunoglobulin production | 0.001 |
| GO:0006355 | regulation of transcription, DNA-templated | 0.008 |
| GO:0006910 | phagocytosis, recognition | 0.035 |
| GO:0000028 | ribosomal small subunit assembly | 0.044 |
| GO:0050853 | B cell receptor signaling pathway | 0.046 |
| GO:0006364 | rRNA processing | 0.086 |
| GO:1904851 | positive regulation of establishment of protein localization to telomere | 0.091 |
| GO:0002181 | cytoplasmic translation | 0.091 |
| GO:0016571 | histone methylation | 0.100 |

Supplementary figures

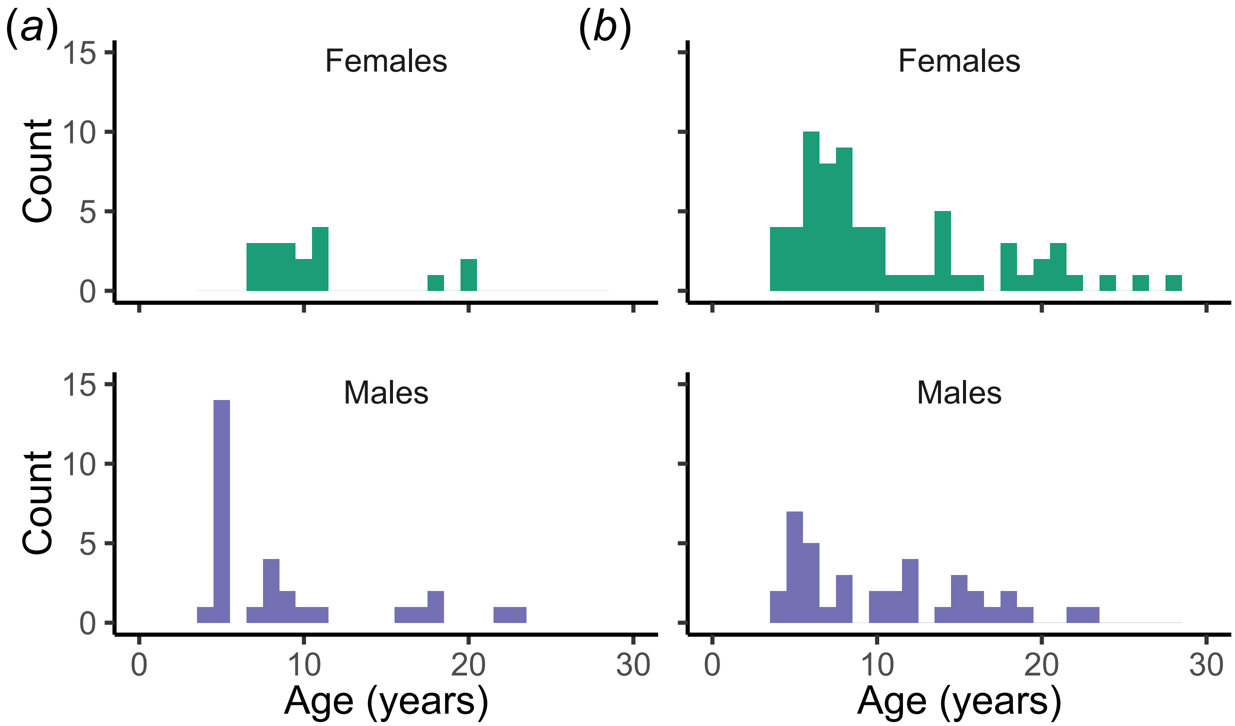


**Figure S1**. Age distributions by sex of (a) the mRNA expression dataset and (b) the CpG methylation dataset presented in this paper.

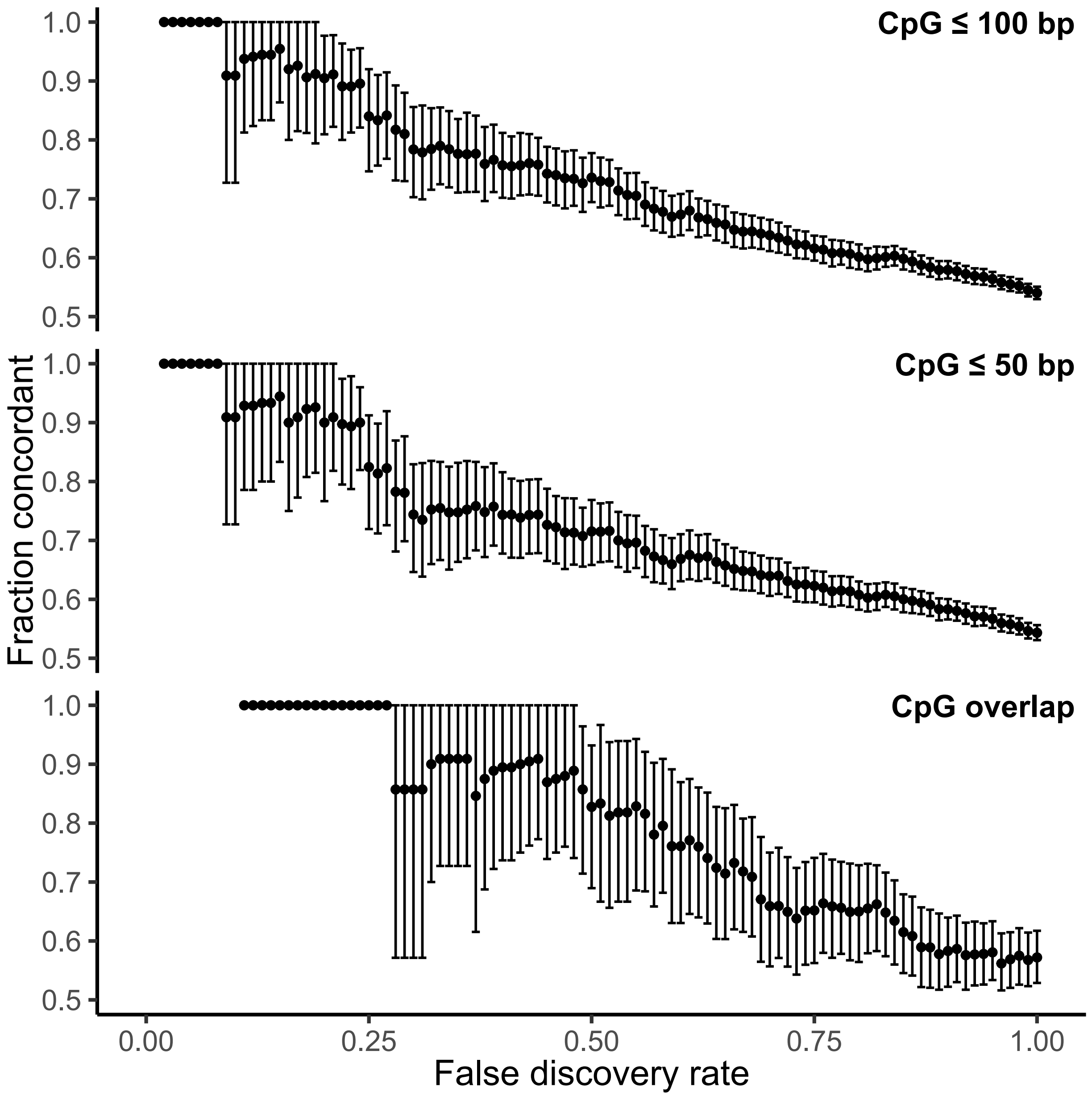


**Figure S2**. Similarities in estimates of concordance were obtained after adjusting criteria for identifying orthologous sites. In the top panel (identical to figure 2d), human CpGs were considered orthologous if they were the nearest neighbors to macaque CpGs and were within 100 bp (4,582 orthologous sites identified with these criteria). In the middle panel, human CpGs were required to be within 50 bp (3,045 orthologous sites identified with these criteria). In the bottom panel, human CpGs were required to overlap completely with macaque CpGs (278 sites).

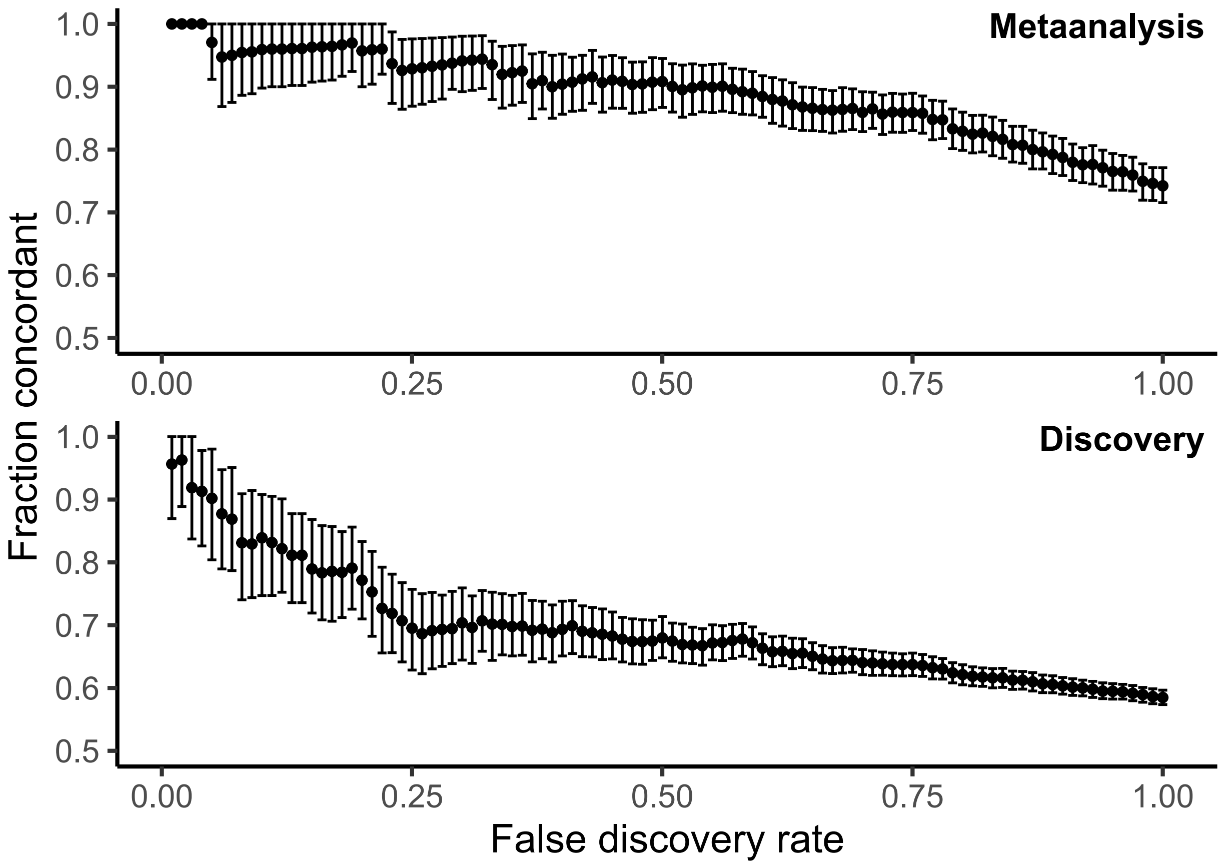


**Figure S3**. Comparison of concordance results obtained when comparing to the separate discovery and metaanalysis results reported by Peters *et al.* [10]. We first compared to results reported from their metaanalysis findings (top panel) because these were corroborated through the use of an additional replication dataset and therefore more strongly supported. When we compared to results reported from their discovery findings (bottom panel), we see lower estimates of concordance throughout, with a steep drop at an FDR threshold of around 0.1.
